# Putting species back on the map: devising a robust method for quantifying the biodiversity impacts of land conversion

**DOI:** 10.1101/447466

**Authors:** América P. Durán, Jonathan M.H. Green, Christopher D. West, Piero Visconti, Neil D. Burgess, Malika Virah-Sawmy, Andrew Balmford

## Abstract

**Aim:** Quantifying connections between the global drivers of habitat loss and biodiversity impact is vital for decision-makers promoting responsible land-use. To that end, biodiversity impact metrics should be able to report linked trends in specific anthropogenic activities and changes in biodiversity state. However, for biodiversity, it is challenging to deliver integrated information on its multiple dimensions (i.e. species richness, endemicity) and keep it practical. Here, we developed a biodiversity footprint indicator that can i) capture the status of different species groups, ii) link biodiversity impact to specific human activities, and iii) be adapted to the most applicable scale for the decision context.

**Location:** Cerrado Biome, Brazil

**Methods:** We illustrate this globally-applicable approach for the case of soybean expansion in the Brazilian Cerrado. Using species-specific habitat suitability models, we assessed the impact of soy expansion and other land uses over 2,000 species of amphibians, birds, mammals and plants for three time periods between 2000 and 2014.

**Results:** Overall, plants suffered the greatest reduction of suitable habitat. However, among endemic and near-endemic species – which face greatest risk of global extinction from habitat conversion in the Cerrado - birds were the most affected group. While planted pastures and cropland expansion were together responsible for most of the absolute biodiversity footprint, soy expansion via direct conversion of natural vegetation had the greatest impact per unit area. The total biodiversity footprint over the period was concentrated in the southern states of Minas Geráis, Goiás and Mato Grosso, but the soy footprint was proportionally higher in those northern states (such as Bahía and Piauí) which belong to the new agricultural frontier.

**Main conclusions:** The ability and flexibility of our approach to examine linkages between biodiversity loss and specific human activities has substantial potential to better characterise the pathways by which habitat loss drivers operate.

## 1 INTRODUCTION

Habitat loss due to land-use change is the biggest threat facing global biodiversity (Gibson et al., 2011; Joppa et al., 2016). Improved quantification of the scale of such change has been essential in supporting international initiatives for better protection and management of land (Ramsar, 1971; CBD, 2002; MEA, 2005). However, further progress is now needed in the identification of underlying drivers of land-use change and the monitoring of their associated environmental impacts to effectively articulate strategies and actions to mitigate them (Han *et al.*, 2014). This need is echoed in the recently approved Sustainable Development Goals (SDGs), particularly in Goals 12 and 15 concerning the sustainable use of land and the responsible production and consumption of its derived commodities.

Biodiversity footprint indicators– which quantify the extent to which human activities impact upon biodiversity – enable the monitoring and reporting of impacts on biodiversity of specific human pressures (Sparks et al., 2011; Hoekstra & Wiedmann, 2014; Hill et al., 2016). Using these to quantify associations between underlying drivers, human activities and biodiversity loss not only helps track the drivers of change (e.g. consumption patterns), but can also reveal the pressures (e.g. agriculture expansion) and mechanisms (e.g. habitat conversion) through which drivers impact biodiversity (Balmford et al., 2009). Biodiversity, however, is a multidimensional concept and comprehensive measurement of changes in its state still poses challenges (Souza et al., 2015). For instance, it could be assessed in terms of species number (Newbold et al., 2015), rarity (Drever, Drever & Sleep, 2012), population density (Collen et al., 2009) and functional diversity (Cadotte, Carscadden & Mirotchnick, 2011). In turn, these dimensions can be variously affected by different mechanisms resulting from human activities, such as habitat loss (Hanski, 2011) or fragmentation (Fahrig, 2003). An increasing understanding of how anthropogenic mechanisms affect different dimensions of biodiversity (Pearson et al., 2014; Pfeifer et al., 2017), together with a larger methodological toolkit (Ewers, Marsh & Wearn, 2010), provides an unprecedented opportunity to standardize a comprehensive biodiversity impact metric. Nevertheless, most commonly-used measuring techniques for the assessment of impacts on biodiversity still focus on change in species richness, which does not capture the whole picture. Furthermore, it is important that when assessing drivers of change, biodiversity impact metrics can translate impact estimates into scales at which information on anthropogenic activities is available and decisions are made (Ewers et al., 2010). When working with relative species richness loss, however, the spatial variability of the impact becomes difficult to scale up as the absence of species identity can lead to challenges such overrepresentation of species’ ranges or misrepresentation of biodiversity priority areas (Veach et al., 2017).

Approaches based on habitat suitability models offer great potential for biodiversity footprint indicators because they can integrate spatially-explicit information on anthropogenic land use and the ecology of individual species (Rondinini et al., 2011; De Baan et al., 2015). Unlike approaches that estimate potential regional or local loss of species richness (Newbold et al., 2015; Chaudhary & Kastner, 2016), models of habitat suitability retain species-specific information Rondinini et al., 2011; de Baan, Mutel, Curran, Hellweg & Koellner et al., 2013), highly relevant given the multiple dimensions of biodiversity. Specifically, they quantify the relative change in the extent of suitable habitat (ESH) arising from land conversion, which allows estimation of a species-specific impact metric that can be associated with a particular human land-use change (Visconti et al., 2011; De Baan et al., 2015). ESH is described by the intersection of a species’ geographic range with its environmental preferences, measured in terms of variables such as vegetation cover, elevation and the location of water bodies and wetlands (Rondinini 138 et al., 2011). Changes in these variables, mainly due to habitat conversion, will reduce the extent of usable habitat, affecting the persistence of local populations (Mantyka-pringle, Martin, Rhodes, 2012).

Previous applications of habitat suitability models have tended to assume that a species’ persistence is directly proportional to the extent of remaining habitat, failing to take into account preceding habitat loss (Buchanan, Donald & Buchart, 2011; van Soesbergen et al., 2017). An important consequence of this for conservation is the underestimation of the impact of current habitat loss on species that have lost a considerable proportion of their original habitat before the assessment is performed (Groves et al., 2002; see section 2.1). Some studies have addressed this limitation but have so far offered limited taxonomic coverage and used projected land-use changes rather than direct observations of habitat conversion (Visconti et al., 2016; Strassburg et al., 2017).

Here, using a non-linear and spatially-explicit approach we describe a biodiversity footprint indicator designed to provide information on biodiversity impact, which can be explicitly linked to specific human activities and adapted to relevant contexts and scales of decision-making. The dual nature of this metric, both footprint and indicator, allows us to quantify the biodiversity impact of specific human activities, while reporting linked trends in pressure (e.g. agriculture expansion), mechanism (e.g. land-use change) and biodiversity state. We illustrate this approach using the example of the cultivation of soybean (*Glycine max*) in the Brazilian Cerrado. We use species-specific habitat suitability models for four taxonomic groups (amphibians, birds, mammals and plants) to illustrate three key benefits of the method: i) its flexibility in capturing and incorporating various levels of ecological information; ii) the scope for linking biodiversity impact to specific human activities such as agricultural commodity expansion; and iii) the capacity to aggregate the estimates across different spatial scales.

## 2 METHODS

An important attribute we have considered during the design of the method is its applicability under a broad range of contexts. We therefore first describe how to implement the approach in general terms (section 2.1), indicating the types of data to use in each step. We then explain how this was implemented, and the specific datasets used, for investigating the impacts of soy expansion in the Brazilian Cerrado (section 2.2).

### 2.1 Method and rational of the biodiversity footprint indicator

The approach involves three steps: i) Mapping the extent of suitable habitat for each species of interest, thus including species’ distributions; ii) Estimating for each species the reduction in their population persistence from the proportional loss of ESH due to land-cover conversion. Within this same step, combining the estimates across species to assess biodiversity impact, thereby considering quantity and variability of species; and iii) Linking biodiversity impact to measures of specific human activities.

#### Mapping the Extent of Suitable Habitat (ESH)

We included all species whose geographic ranges intersect the study region for which habitat information is available. Every grid cell in a species’ range in the region can be coded as suitable if both the following conditions are met: (i) the cell is within the geographic range of the species, and (ii) the local environment is within the species’ known preferences (in terms of land cover, elevation, etc.). The latter requires the harmonisation, in consultation with experts, of the categories of the available land-cover map with those used to describe species’ habitat preferences. Coding of the suitability of cells should be repeated in the same way for various points in time using environmental data appropriate for each time. Each time period at which suitability is determined should be assessed against a benchmark time. For migratory species, ESH should be mapped separately for each species’ resident, breeding and non-breeding ranges, based on seasonal differences in their habitat preferences. This accounts for seasonal variation in species’ habitat requirements.

#### Estimating the marginal value of suitable habitat

The next step involves calculating, for each species, the remaining proportion of its initial benchmark ESH within the study area at each subsequent point in time. Changes in ESH are then used to derive a non-linear persistence score, *P*, which captures the cumulative effect of habitat loss on the likelihood of the species’ persistence in the study region:

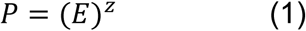

where *E* is the remaining proportion of the original ESH, and *z* is the extinction coefficient. Equation (1) is analogous (at the level of a single species; Thomas et al., 2004) to the community-level species-area curve (*S*=*cA^z^*). We propose its use here based on the conjecture that the conversion of given absolute area of suitable habitat (*A_loss_* in Fig. 1) after a species has lost a small amount of its initial benchmark ESH (*a* in Fig. 1), is likely to reduce the probability of the species’ persistence less (*c* in Fig. 1) than if the same area of suitable habitat was lost after much of the initial ESH had already been converted (*b* and *d* in Fig. 1). As an increasing number of studies have demonstrated, historical habitat loss can have important cumulative and delayed effects on biodiversity (Krauss et al., 2010; Wearn et al., 2012), and ignoring such effects by assuming, for example, a linear relationship between habitat loss and species’ persistence (equivalent to a z-value of 1 in Equation 1), can result in sever underestimations of biodiversity loss.

**Figure 1.**
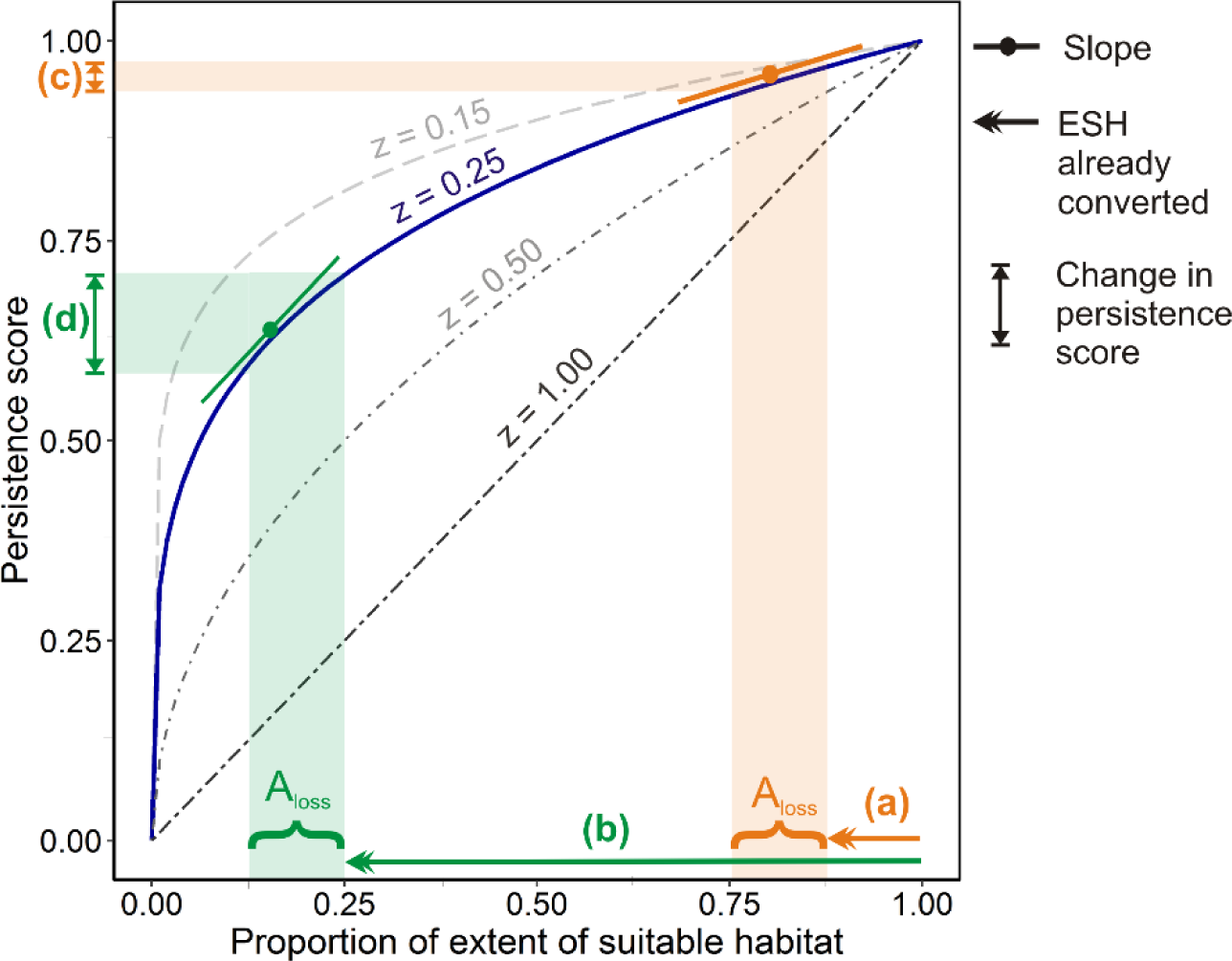
Relationship between remaining extent of suitable habitat and species’ persistence score, P, upon which the biodiversity footprint is based. If this relationship follows a power law (with 0<z<1) the loss of a given area of suitable habitat (A_loss_), when only a small proportion (a) of the original habitat has been lost previously, has a smaller impact (c) on score P than losing the same area when a much larger proportion (b) has already been lost (d). The size of this difference depends on the extinction coefficient z, which may well vary across taxa and regions.

In addition, distribution size (here estimated by ESH), is one key factor contributing to extinction risk and it is also closely correlated with population size (Blackburn et al., 1997; Harris & Pimm, 2008). Therefore, reduction in species distribution is expected to affect populations’ persistence (IUCN, 2001). Here we used proportion of ESH (and not the absolute area), as this allows assessing impact across species in a standardized way while accounting for quantity and variability of species. Since the initial benchmark ESH reflects the historical distribution size (when probability of persistence was 1; see below for more details), the proportional area loss declines at the same rate that absolute area loss relative to the benchmark. Consequently, species with restricted ranges will move faster to the left along the curve as the loss of one unit of absolute area means a higher proportional loss than for a widespread species. It is worth noting, however, that further work is required to establish empirically how the absolute and proportional area losses of individual species are related to probability of persistence. As yet, there is no standard method for such a calculation.

Once *P* has been estimated for two or more time points, the effect of intervening habitat loss on a species’ likelihood of persistence within the study area can be calculated as Δ*P*, the corresponding difference in *P*-values:

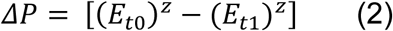

where *E*_*t*0_ and *E*_*t*1_ are the remaining proportions of ESH at *t*_0_ and *t*_1_, respectively.

For migratory species, an overall Δ*P_mig_* score should be calculated from Δ*P_mig_* scores derived separately for the species’ breeding and non-breeding ESH. In order to estimate the total change in a migratory species’ persistence score, a multiplicative effect can be assumed, as previously suggested by empirical (Lockwood, 2004) and theoretical studies (Iwamura et al., 2013):

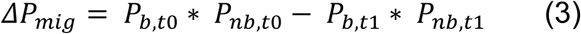

where *p_b_* and *p_nb_* are the persistence scores within the breeding and non-breeding ranges, respectively. This approach accounts for an interactive effect between populations’ likelihood of persistence along the migratory movements - an important effect to consider in biodiversity impact quantifications (see Appendix S1 and Fig. S1.1 in Supporting Information for further discussion of the implications of this approach).

If there is interest in estimating global-level impacts but the study region itself is not global, each species’ Δ*P*-values should be weighted by the proportion of its global geographic range falling within the study region. Other ways of weighting different species – to reflect their ecological or evolutionary significance, for example – can also be employed at this stage (see Discussion). The weighted Δ*P* of each species (including migratory ones) will then be assigned to individual cells to derive the marginal value of the loss of suitable habitat, *MV*, for each cell converted over a given time interval. Thus, for a period of time *t*_0_→*t*_1_, the marginal value of the loss of suitable habitat within cell *j* (belonging to a set of converted cells *R*), for the weighted Δ*P* score of species *k*, *MV*_*t*0-*t*1,*j, k*_, can be represented as:

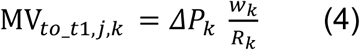

where *R* is the total number of cells converted from suitable to unsuitable for that species in the period *t*_0_→*t*_1_, and *w* is the weight of species *k* (representing, for example, the proportion of its geographic range falling within the study region). The 272 resulting distribution maps of marginal loss values for individual species are then overlaid and values summed across species to obtain, for each cell, an aggregated biodiversity impact metric. Using maps of administrative boundaries (e.g. municipalities), the cell-level impact values can then be aggregated to give totals for administrative units of interest.

#### Linking biodiversity impact to a human pressure

The biodiversity impact scores described above can be attributed to categories of land-use conversion due to different types of human use (e.g. natural vegetation to cropland). Where more detailed spatial information on human activities is available, the impacts can be associated even more specifically with particular production systems, which in turn are related to the land-use conversion assessed above (Eq. 2).

### 2.2 Applying the method to soy expansion in the Cerrado

We applied the approach outlined above to the specific case of the expansion of soy cultivation in the Cerrado over the period 2000 – 2014. Considered one of the world’s most diverse savannah ecosystem, the Cerrado is severely threatened by the expansion of soybean cultivation and cattle ranching (Strassburg et al., 2017).

#### Mapping ESH within the biome

After selecting all the species in our focal taxa whose current ranges intersected the Cerrado boundary (IBGE 2004) and for which habitat information was available, we produced habitat suitability models to obtain 234 ESHs for amphibians, 846 for birds and 288 for mammal species, each at 250 m x 250 m resolution (the resolution of the best available land cover maps for Brazil with which land-use change can be quantified consistently; IBGE 2015). Based on information on habitat associations and elevation limits obtained from the IUCN Habitats Classification Scheme (IUCN 2017), we refined the historical geographic range (Extant, Probably Extant, Possibly Extinct, Extinct and Presence Uncertain) of each vertebrate species (BirdLife 2016, IUCN 2017) using a digital elevation model (USGS 2006) and land cover maps (IBGE 2004, 2011, 2014). The 14 categories of the land-cover map were harmonised with the 74 habitat preference levels (for details see Appendix S2 and Table S2.1 in Supporting Information).

Multiple environmental variables define species distribution as well as a populations’ response to habitat loss. Yet, data on habitat preferences and altitudinal range are the only species-specific variables that are available globally. Since this approach aims to be globally applicable we limited the illustration of ESH mapping to these two variables, although further information can be incorporated during the refinements of species ranges (see Discussion).

We also produced habitat suitability model maps for 648 plant species whose ranges intersect the Cerrado. In the absence of more detailed information on species’ habitat requirements, we refined their geographic ranges (Martinelli & Moraes 2013) using information on vegetation types. We assumed that only those vegetation categories classified as natural by the Brazilian Institute of Geography and Statistics (IBGE) were potentially suitable for these species, while other semi-and non-natural categories were unsuitable (Table S2).

We applied our approach to IBGE land cover maps for the year 2000, 2010, 2012 and 2014. For vertebrates, we used a map of original vegetation cover for the Cerrado as our initial benchmark (c.a. 16th century; IBGE 2004), to estimate ESH prior to large-scale cultivation for each species. For plants, the geographic range intersecting the study region was considered to delineate its original ESH.

#### Impacts of habitat loss on the Cerrado’s biodiversity

For the calculation of weighted marginal values of each cell for each species (Eq. 4), we adopted a *z*-value of 0.25, based upon its ability to predict proportions of species becoming extinct or threatened as a result of habitat loss in several species-area analyses (Brooks & Balmford, 1996; Brooks, Pimm & Oyugy et al., 1999). Different *z*-values influence the effect of habitat loss on probability of persistence (Fig. 1; Eq. 2). However, our qualitative conclusions concerning the relative role of human activities on different groups of species and the spatial distribution of estimated biodiversity impacts are not strongly dependent on the choice of a particular value of *z* (Table S3.2 and Fig. S3.2 in Appendix 3 for the effects of plausible variation in *z*-coefficients). In addition, when information is available different z-values can be assigned to different biodiversity groups. Species’ Δ*P* values were weighted by the proportion of their global geographic range falling within the study region.

We also considered marginal increases due to gain of suitable cells (e.g. through reversion of converted land to natural habitat). However, reversion is currently on such a limited scale in the Cerrado that incorporating such gains had a minor impact on the results for most of the groups, and was therefore not considered in the main text (see Fig. S4.3 in Appendix S4).

#### Quantifying the biodiversity impact of soybean expansion

We used two types of maps for cumulative soy expansion from Gibbs et al. (2015) for the period 2000-2014: 1) direct expansion of soy into natural vegetation (where soy production occurred within three years of natural vegetation conversion); and 2) expansion into previously-cleared areas. Before intersecting soy-expansion maps with biodiversity impact maps, we combined the former with IBGE land-conversion maps to distinguish soy expansion from non-soy crop expansion (see Fig. S5.4 in Appendix S5 in Supporting Information for details on land-conversion maps analysis). The resulting merged layer allowed us to also assess the impact of other non-crop categories such as planted pasture.

## 3 RESULTS

### 3.1 Assessing the biodiversity footprint for different species groups

In order to illustrate biodiversity impact at species level we focused on five conservation flagship species in the Cerrado (WWF, 2015). For the Maned Wolf (*Chrysocyon brachyurus*), Jaguar (*Panthera onca*), Giant Armadillo (*Priodontes maximus*), South American Tapir (*Tapirus terrestris*) and Giant Anteater (*Myrmecophaga tridactyla*), habitat loss within the Cerrado has caused steady declines in their weighted persistence scores over the 2000-2014 period (Fig. 2a; declines of 0.006, 0.007, 0.009, 0.009, and 0.01, respectively). As the only species for which ‘Arable’ and ‘Pasture’ are considered suitable habitats (IUCN 2017), the Maned Wolf presented the smallest decline of the five species and had a markedly higher score than the other species in 2014. Major losses of natural vegetation had occurred by 2000 already, with Giant Armadillo losing 80% of its Cerrado ESH, Giant Anteater 83%, Jaguar 88% and South American Tapir 84%. While the Jaguar showed the largest reduction of its original ESH within the Cerrado, this accounts for a relatively small proportion of its global range, resulting in a smaller change in its persistence score than for other species (Fig. 2a).

**Figure 2.**
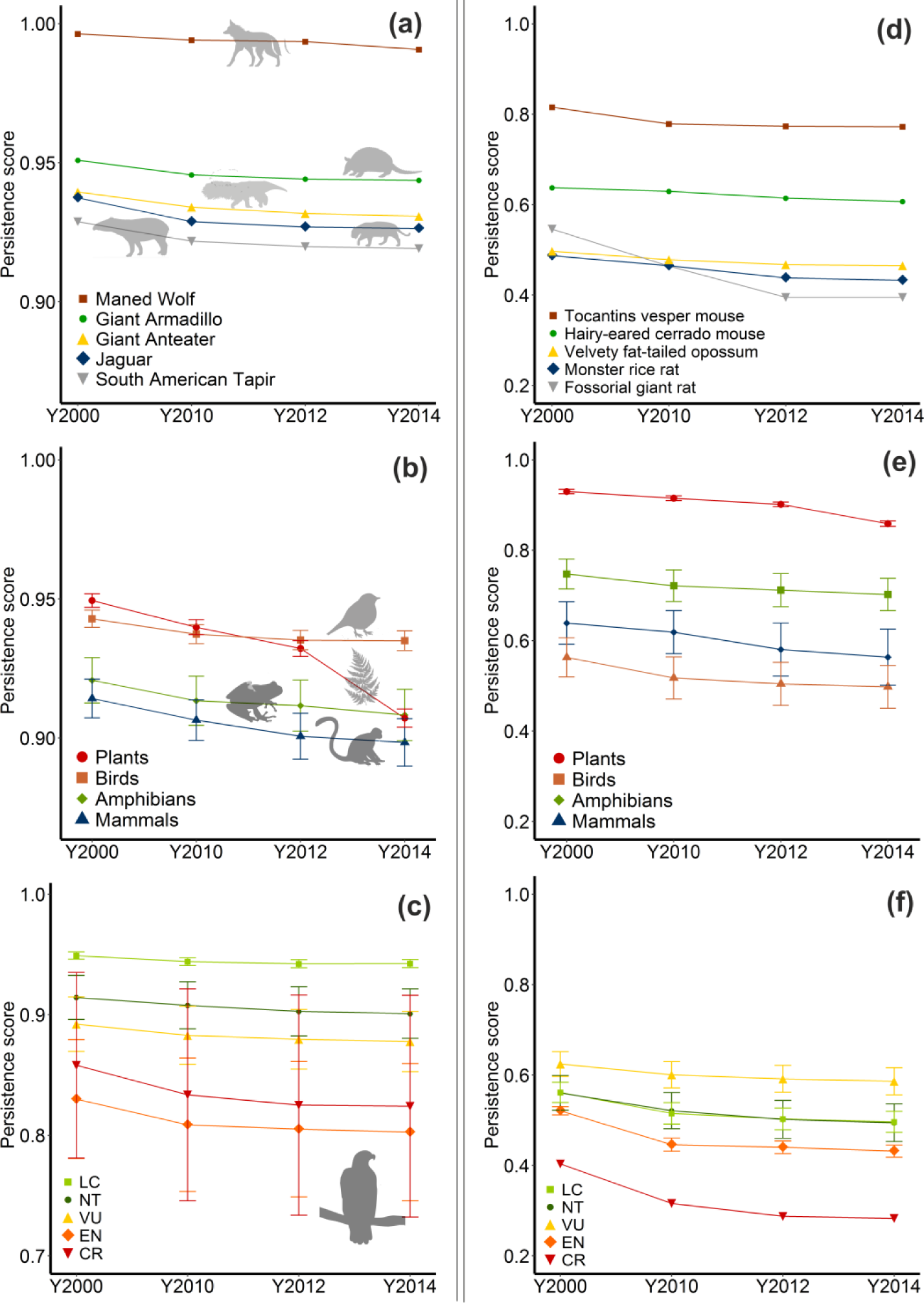
Changes in persistence score due to land conversion between 2000 and 2014, calculated for different levels and elements of biodiversity: (a) at species level, showing results for five flagship species; (b) for taxonomic groups, showing mean persistence scores for four vertebrate taxa; (c) grouped by IUCN Red List status, showing mean persistence scores for birds; (d) at species level, showing results for five endemic and near-endemic mammal species; (e) for taxonomic groups, showing mean persistence scores for endemic and near-endemic species only; and (f) grouped by IUCN Red List status, showing mean persistence scores for endemic and near-endemic bird species only. Upper and lower bars show one standard error.

When biodiversity impacts were aggregated by taxonomic group (Fig. 2b), plants showed the largest impact 2000-14 (0.042 ± 0.002; mean ± standard error), then mammals (0.015 ± 0.007), amphibians (0.012 ± 0.008) and birds (0.0079 ± 0.003). In the 2012-2014 period alone, plants lost on average 9.1% of their original ESH within the Cerrado (0.30 y^−1^), compared to the 7.1% lost over the 2000-2012 period (0.07 y^−1^). This resulted in a sharper mean decline of plants’ weighted persistence score (0.025 ± 0.0032; 0.047/y^−1^), relative to the prior twelve years (0.017 ± 0.0013; 0.006/y^−1^).

Focusing on birds, we also assessed how impacts varied across species of different conservation status (Fig. 2c). Declines in persistence scores were consistently greater among species in higher extinction risk categories (Fig. 2c). Among Critically Endangered (CR; 0.034 ± 0.015) and Endangered (EN; 0.027 ± 0.007) species, the mean persistence score decreased more between 2000 and 2014 than among Vulnerable (VU; 0.014 ± 0.002), Near Threatened (NT; 0.013 ± 0.002) and Least Concern (LC; 0.006 ± 0.0004) species.

We also assessed endemic and near-endemic species, for which we included those species with more than 70% of their global range falling within the Cerrado. Overall, compared to more widely distributed species and species groups (Fig. 2a,b,c), endemics presented a much more severe decline in their persistence score (Fig. 2d,e,f), representing an acute threat to their global persistence.

### 3.2 Links between biodiversity footprint and commodity production

For 2000-2014, our results revealed that conversion to grassland, whilst comprising 45% of the area of habitat converted, was responsible for just 14% of the total biodiversity footprint of all land conversion in the region (Fig. 3). In contrast, planted pastures, crops other than soybean and mosaic crops were together responsible for 43% of the habitat conversion but 67% of the biodiversity footprint (Fig. 3). Soybean expansion into previously converted habitat was responsible for 3% of the habitat conversion but 5% of the total biodiversity footprint. Lastly, whilst direct expansion of soy into natural vegetation was responsible for only 0.15% of the total habitat converted, it accounted for 0.8% of the biodiversity footprint.

**Figure 3.**
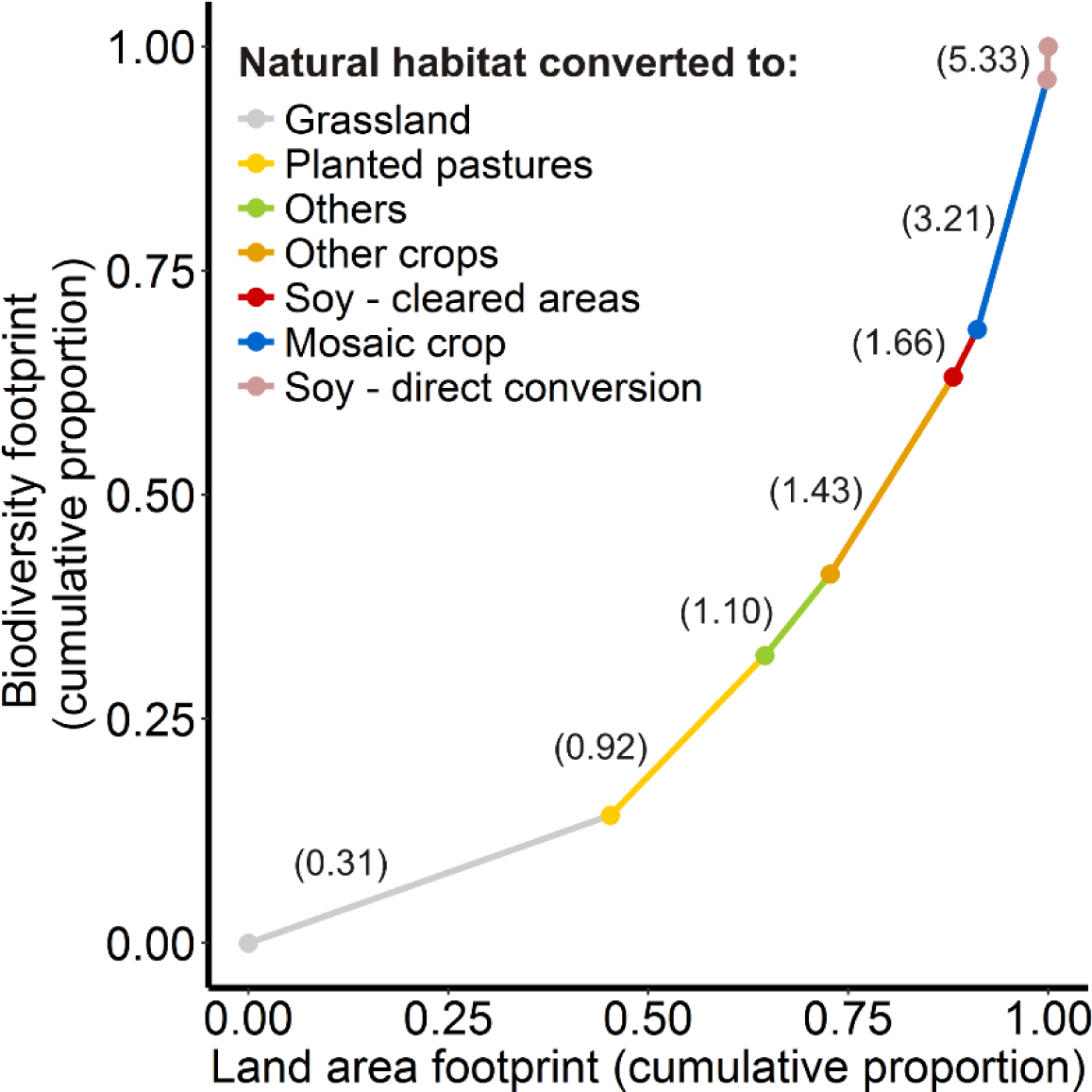
The proportional contribution of different land-use conversions to the total biodiversity footprint 2000-14 in the Cerrado including four taxonomic groups, plotted against the proportion of the total land area footprint of each land-use change. Land-use conversions are plotted in order of increasing ratio of proportional contribution to change in persistence score: proportional contribution to loss of ESH (with the ratios shown in parentheses). Higher ratios thus indicate land-use conversions with disproportionately high impacts on our biodiversity footprint metric given the area converted. We aggregated IBGE land-use categories as follows: other crop (than soybean), planted pasture, mosaic (mosaic-forest, mosaic-crop and mosaic shrubland), grassland, and other.

We also explored the relative footprint per unit area, which can reveal land use transitions with disproportionate impacts on biodiversity. To this end, we calculated the ratio of proportional contribution to the total biodiversity footprint to proportion of area converted (Fig. 3): higher ratios indicate land-use conversions with disproportionately high biodiversity footprints. We found that, although soy expansion through direct conversion of natural habitat had the smallest areal footprint, it had the highest impact on biodiversity per unit area (5.3). This indicates that soy expansion through direct conversion has altered a disproportional amount of ESH relative to the total footprint land used. Further disaggregating this by taxa showed that mammals were the most affected group (8.3), followed by birds (7.9), amphibians (2.6) and plants (1.6) (see Fig. S6.5 in Supporting Information).

### 3.4 Adapting biodiversity footprint to scales of decision-making

We designed our footprint indicator so it can be aggregated at different scales, while still capturing ecological impacts of change. Each cell’s score contributes proportionally to the footprint (Eq. 4), so cell values can be summed across any area of interest (e.g. a municipality) to reflect that area’s contribution to the overall footprint. In the Cerrado, aggregating the biodiversity footprint indicator across municipalities and states for the 2000-14 time period revealed distinct insights at different scales (Fig. 4a-c). It is possible to identify municipalities with relatively high biodiversity impact within states of relatively low footprint, revealing local-scale impacts that are diluted at coarser resolution. For example, the municipalities of Mateiros (with a score of 0.54) and Jaborandi (0.32), fell in two states with overall low values: the state of Tocantins (1.36) and Bahía (0.99), respectively. These two states are part of the ‘MATOPIBA’ agricultural frontier and have undergone more rapid habitat conversion since 2000 than other states (Fig. S7.6 in Supplementary Information). Their relatively low species richness (Fig. S8.7b), however, results in lower overall impact scores than in more biodiversity-rich states (Fig. S8.7b; Fig. S9.8). Nevertheless our method singles-out areas that may be of particular conservation concern in these states. Our method also allowed us to disentangle the state-level biodiversity footprint into different types of land conversion (Fig. 4d). In two states that underwent particularly extensive habitat clearance prior to 2000, Mato Grosso and Goiás, subsequent soy expansion was largely into already-cleared areas (Fig. 4d), and so had a relatively low footprint. In contrast, in Bahía and Piauí, two states that have undergone extensive habitat clearance within the new agricultural frontier, recent soy expansion is associated with a greater impact on biodiversity.

**Figure 4.**
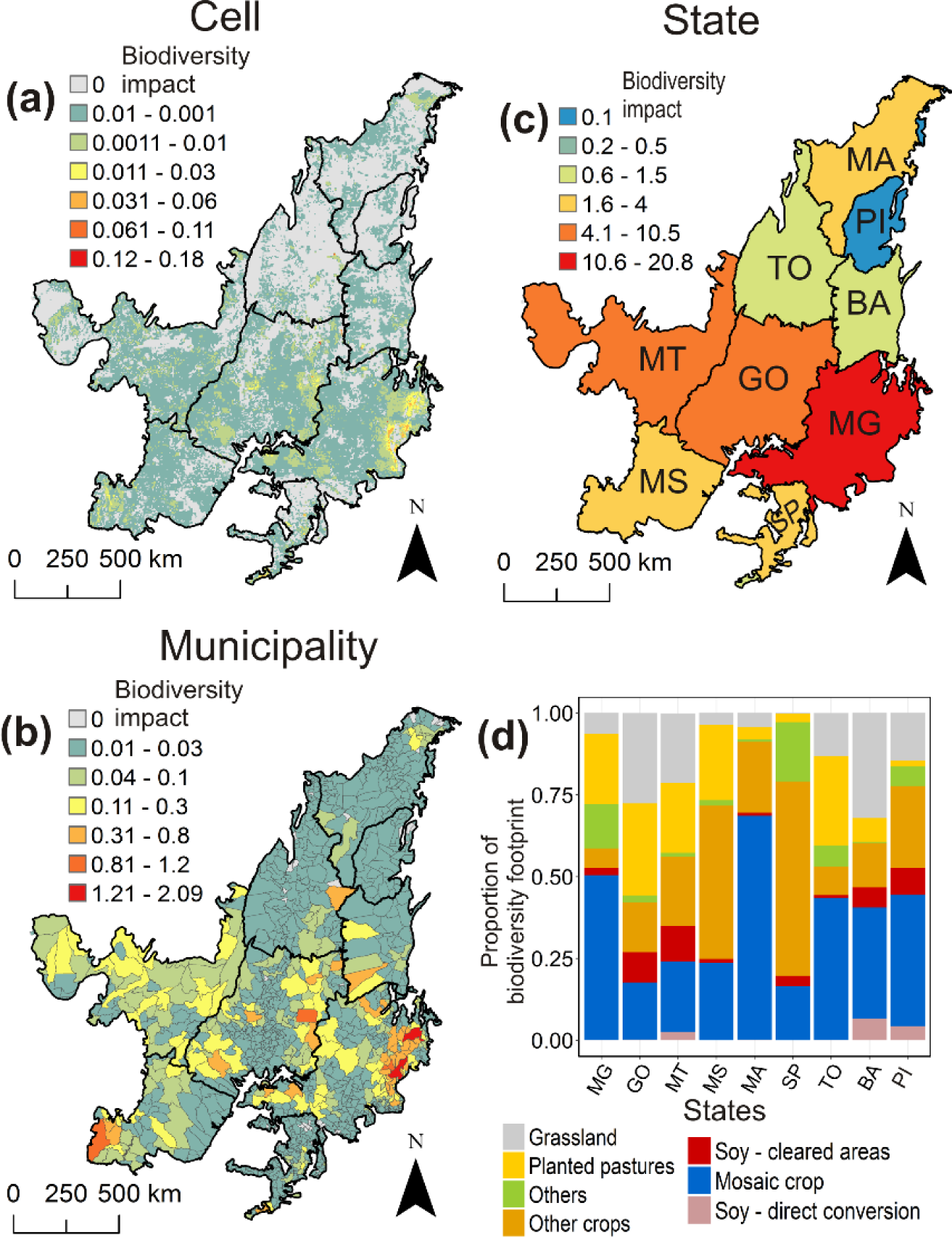
Distribution of biodiversity footprint scores due to loss of ESH 2000-14, when data are aggregated for all species at three different spatial scales and for different land/use changes. (a) Cell (0.0625 km^2^); (b) Municipality; (c) State; and (d) proportional contribution of different land-use conversions to the total biodiversity footprint 2000-14 at state level. [MG: Minas Geráis; GO: Goiás; MT: Mato Grosso; MS: Mato Grosso do Sul; MA: Maranhão; SP: São Paulo; TO: Tocantins; BA: Bahia; PI: Piauí].

## 4 DISCUSSION

Three criteria shaped the design of our biodiversity indicator; it should: i) be able to capture the status of different components of biodiversity, while including aspects such as the distribution, quantity and variability of species, ii) allow biodiversity impacts to be linked to specific human activities; and iii) be scalable to inform decision-making at different levels. Below we highlight strengths and limitations of our approach in relation to these criteria.

### 4.1 Capturing the status of different components of biodiversity

By combining information on land cover change, individual species’ distributions and habitat preferences this method identifies which biodiversity elements are most affected and where the greatest impacts have occurred (Fig. 2; Fig. 4). Working at the level of species allows features of the ecology of species (such as their habitat specificity, endemicity or migratory movements) to be incorporated, thereby considering species’ distribution and variability (Visconti et al., 2016). Making such information spatially explicit, allows hotspots of biodiversity risk to be identified, and provides information on the quantity of species that are vulnerable (Visconti et al., 2011).

When information on habitat preferences is unavailable, as it was here for plants, assumptions on habitat requirements need to be made. If such assumptions are generous – such as that species can occur in a wide range of land covers including anthropogenic ones - there is higher chance of incurring errors of commission (assuming a species occurs where it does not) and hence of underestimating species’ risk of local extinction (Rondinini, Stuart & Boitani, 2005). In contrast, more conservative assumptions are prone to errors of omission (incorrectly assuming that a species is absent) and thus of overestimating impact (Rondinini et al., 2005). Under the precautionary principle, widely adopted in assessing biodiversity risk (Myers 1993; Dickson & Cooney, 2005), conservative assumptions might be more appropriate.

As our study area covers a fraction of the global population of many species, we weighted species’ persistence scores by the proportion of their geographic range that intersects the Cerrado. This assigns more weight to impacts on those species restricted to the biome, but places less emphasis on the local loss of species with a small fraction of their range intersecting the Cerrado. While such losses might have limited global conservation consequences, they could nonetheless have significant ecological or cultural effects. For instance the Jaguar experienced only a small change in its weighted persistence score as a result of habitat loss in the Cerrado; a non-weighted score shows much more extensive decline (a local decline to 0.51 versus a global decline to 0.92 by 2014; Fig. S10.9a). To represent losses of culturally- or ecologically-important species, it would also be possible to apply additional weightings when summing Δ*P* values across species, which could reflect variation in ecological or cultural significance.

While the wide availability of the data used here makes our method practical and accessible, we acknowledge that the variables we use cannot fully capture the ecological complexity to which species respond. For instance, habitat fragmentation and isolation can be important determinants of species occurrence (Ewers et al., 2010) and ignoring such landscape-level information can add further error into species’ distribution mapping (i.e. omission and commission error, see above for further discussion). Even though information on how species respond to fragmentation and edge-effects is currently absent from the IUCN Red List, recent studies have provided insight in how best to model this. By combining suitable habitat modelling techniques and spatial layers, a continuous representation of individual species’ responses to fragmentation and edge-effects can be calculated (Ewers et al., 2010; Pfeifer et al., 2017). Thus, combining these layers with a biodiversity footprint metric, such as the one proposed in this paper, can help us understand how biodiversity responds to changes in both landscape composition and structure. Such an advancement will provide key insights into land management and biodiversity conservation.

### 4.2 Linking biodiversity impact to specific human activities

Our method disentangles, at different spatial scales, the effects of human activities bringing about habitat loss (Fig. 3). This is essential for then tracking the pathway through which underlying drivers of habitat loss operate (Moran & Kanemoto, 2017). In this study we focused on soy production as the direct human activity affecting habitat loss, which in turn can be influenced by remote drivers such as consumption patterns (de Ruiter et al., 2017), production shortages (Godfray et al., 2010) and population growth (Dasgupta & Ehrlich, 2013). As well as remote drivers, other set of indirect channels can also influence the effects of a human activity on habitat loss. This can be through land use displacement, a widely recognized mechanism underlying indirect land use change (Lambin & Meyfroidt, 2011). Also, via the ability to influence regional land markets, therefore affecting deforestation decisions indirectly (Richards, 2015). Similar to previous studies (Richards, 2015), our estimation of soy indirect impact (through the displacement of cattle ranches into natural vegetation (see Appendix 11 and Fig.11.10 in Supplementary Information)), also suggested a limited role of land use displacement in the overall impact. Thus, incorporating multiple techniques to capture direct and indirect drivers, while encompassing a broader time frame that allows assessing historical land-conversion trends, will certainly better capture the full responsibility of assessed human activities, such as the case of soy in the Cerrado.

### 4.3 Aggregating biodiversity impact at different spatial scales

Developing tools that capture and translate the ecological scale of the problem to scales where decisions are made has been suggested as a key solution to improve evidence impact (Guerrero, McAllister, Corcoran & Wilson, 2013). The results presented here suggest that our proposed method meets these requirements, by capturing relevant ecological information such as species richness, mean historical habitat losses and endemicity (Fig. S8.7), which can be adapted to different scales of decision-making. Metrics of impact that are adaptable to different scales of threat information are also likely to be useful in evaluating causal connections between biodiversity impact and human activities (see section 4.4 for more discussion on this regard). Another relevant aspect is the sensitivity of the aggregated metric to its key parameters (Eq. 2). Using different z-values we observed only minor changes in the aggregated biodiversity footprint and the distribution of biodiversity risk hotspots (Fig. S3.2e,f). As z increases, the decline of species’ persistence score increases for a given loss of ESH (Fig. 1). Hence at higher z, areas (e.g. states) that harbour species with high historical ESH loss such as Mato Grosso (MT) and Mato Grosso do Sul (MS) (Fig. S8.7c), have a higher increment in their aggregate biodiversity footprint than do areas with less historical loss of ESH.

### 4.4 Implications of our method for the Cerrado

The Cerrado example illustrates how our approach can quantify human activities driving land-use change and monitor their biodiversity impacts. Although these activities are well known to be in the Cerrado soy and livestock production, there remains a clear need to map the underlying trade system of both commodities (Garrett, Lambin & Naylor, 2013). Brazil is now the second-largest soy producer worldwide, and in 2013/2014 about half (52%) of soybeans produced in Brazil came from the Cerrado (INPUT, 2016). A better understanding of the highly complex production-to-consumption system, comprising large numbers of trade actors (e.g. producers, manufacturers, exporters), is an ongoing and challenging effort (Godar, Suavet, Gardner, Dawkins, & Meyfroidt, 2016). By linking spatially-explicit biodiversity risk hotspots with information on soy and livestock production and trade our approach provides a platform to start disentangling the relative roles of different actors.

## Data Accessibility

All data used in this study are freely available online (Please see references for more details). Species ranges were obtained from: (https://www.iucnredlist.org/resources/spatial-data-download-Mammals&Amphibians), (http://datazone.birdlife.org/species/requestdis-Birds), and (https://tinyurl.com/y7zxzxhv-Plants). Habitat preferences for vertebrates, including altitudinal ranges, we obtained from (http://apiv3.iucnredlist.org/api/v3/docs). Land cover and boundary data for Brazil were obtained from (https://www.lapig.iesa.ufg.br/lapig/). Digital elevation data was obtained from (https://lta.cr.usgs.gov/products_overview). Soy expansion data were obtained from (Gibbs et al., 2015).

## REFERENCES

De Baan, L., Mutel, C. L., Curran, M., Hellweg, S., & Koellner, T. (2013). Land use in life cycle assessment: global characterization factors based on regional and global potential species extinction. Environmental science & Technology, 47, 9281–9290.

de Baan, L., Curran, M., Rondinini, C., Visconti, P., Hellweg, S., & Koellner, T. (2015). High-resolution assessment of land use impacts on biodiversity in life cycle assessment using species habitat suitability models. Environmental science & Technology, 49, 2237–2244.

Balmford, A., Carey, P., Kapos, V., Manica, A., Rodrigues, A. S., Scharlemann, J. P., & Green, R. E. (2009). Capturing the many dimensions of threat: comment on Salafsky et al. Conservation Biology, 23, 482–487.

BirdLife International and NatureServe (2016) Bird species distribution maps of the world. BirdLife International, Cambridge, UK and NatureServe, Arlington, USA.

Blackburn, T.M., Gaston, K.J., Quinn, R.M., Arnold, H. & Gregory, R.D. (1997) Of mice and wrens: the relation between abundance and geographic range size in British mammals and birds. Philosophical Transactions of the Royal Society of London B: Biological Sciences, 352, 419–427.

Brooks, T., & Balmford, A. (1996). Atlantic forest extinctions. Nature, 380, 115.

Brooks, T. M., Pimm, S. L., & Oyugi, J. O. (1999). Time lag between deforestation and bird extinction in tropical forest fragments. Conservation Biology, 13, 1140–1150.

Buchanan, G. M., Donald, P. F., & Butchart, S. H. (2011). Identifying priority areas for conservation: a global assessment for forest-dependent birds. PloS one, 6, e29080.

CBD (2002) Rio+20 United Nations Conference on Sustainable Development.

Cadotte, M. W., Carscadden, K., & Mirotchnick, N. (2011). Beyond species: functional diversity and the maintenance of ecological processes and services. Journal of Applied Ecology, 48, 1079–1087.

Chaudhary, A., & Kastner, T. (2016). Land use biodiversity impacts embodied in international food trade. Global Environmental Change, 38, 195–204.

Collen, B., Loh, J., Whitmee, S., McRAE, L., Amin, R., & Baillie, J. E. (2009). Monitoring change in vertebrate abundance: the Living Planet Index. Conservation Biology, 23, 317–327.

Dasgupta, P. S., & Ehrlich, P. R. (2013). Pervasive externalities at the population, consumption, and environment nexus. Science, 340(6130), 324–328.

Dickson, B. & Cooney, R. (2005). Biodiversity and the precautionary principle: risk and uncertainty in conservation and sustainable use. London, Earthscan.

Espírito-Santo, M. M., Leite, M. E., Silva, J. O., Barbosa, R. S., Rocha, A. M., Anaya, F. C., & Dupin, M. G. (2016). Understanding patterns of land-cover change in the Brazilian Cerrado from 2000 to 2015. Philosophical Transactions of the Royal Society B, 371, 20150435.

Ewers, R. M., Marsh, C. J., & Wearn, O. R. (2010). Making statistics biologically relevant in fragmented landscapes. Trends in Ecology & Evolution, 25, 699–704.

Fahrig, L. (2003). Effects of habitat fragmentation on biodiversity. Annual Review of Ecology, Evolution, and Systematics, 34, 487–515.

Garrett, R. D., Lambin, E. F., & Naylor, R. L. (2013). Land institutions and supply chain configurations as determinants of soybean planted area and yields in Brazil. Land Use Policy, 31, 385–396.

Gibbs, H. K., Rausch, L., Munger, J., Schelly, I., Morton, D. C., Noojipady, P.,…Walker, N.F. (2015). Brazil’s soy moratorium. Science, 347, 377–378.

Gibson, L., Lee, T. M., Koh, L. P., Brook, B. W., Gardner, T. A., Barlow, J.,…Sodhi, N. S. (2011). Primary forests are irreplaceable for sustaining tropical biodiversity. Nature, 478, 378.

Godar, J., Suavet, C., Gardner, T. A., Dawkins, E., & Meyfroidt, P. (2016). Balancing detail and scale in assessing transparency to improve the governance of agricultural commodity supply chains. Environmental Research Letters, 11, 035015.

Godfray, H. C. J., Beddington, J. R., Crute, I. R., Haddad, L., Lawrence, D., Muir, J. F.,… Toulmin, C. (2010). Food security: the challenge of feeding 9 billion people. Science, 327, 812–818.

Groves, C. R., Jensen, D. B., Valutis, L. L., Redford, K. H., Shaffer, M. L., Scott, J. M.,…Anderson, M. G. (2002). Planning for Biodiversity Conservation: Putting Conservation Science into Practice: A seven-step framework for developing regional plans to conserve biological diversity, based upon principles of conservation biology and ecology, is being used extensively by the nature conservancy to identify priority areas for conservation. AIBS Bulletin, 52, 499–512.

Guerrero, A. M., McAllister, R. Y. A. N., Corcoran, J., & Wilson, K. A. (2013). Scale mismatches, conservation planning, and the value of social-network analyses. Conservation Biology, 27, 35–44.

Han, X., Smyth, R. L., Young, B. E., Brooks, T. M., de Lozada, A. S., Bubb, P.,…& Turner, W. R. (2014). A biodiversity indicators dashboard: Addressing challenges to monitoring progress towards the Aichi biodiversity targets using disaggregated global data. PloS one, 9, e112046.

Hanski, I. (2011). Habitat loss, the dynamics of biodiversity, and a perspective on conservation. AMBIO: A Journal of the Human Environment, 40, 248–255.

Harris, G., Pimm, S.L. (2008) Range size and extinction risk in forest birds. Conservation Biology, 22, 163–171.

Hill, S. L., Harfoot, M., Purvis, A., Purves, D. W., Collen, B., Newbold, T.,…. Mace, G. M. (2016). Reconciling biodiversity indicators to guide understanding and action. Conservation Letters, 9, 405–412.

Hoekstra, A. Y., & Wiedmann, T. O. (2014). Humanity’s unsustainable environmental footprint. Science, 344, 1114–1117.

Input (2016) The expansion of soybean production in the Cerrado. Available:http://www.inputbrasil.org/wp-content/uploads/2016/11/The-expansion-of-soybean-production-in-the-Cerrado_Agroicone_INPUT.pdf. Accessed: January 2018.

Instituto Brasileiro de Geografia e Estatística (IBGE). 2004. Borders of Brazilian biomes. http://maps.lapig.iesa.ufg.br/lapig.html. Accessed August 2016.

Instituto Brasileiro de Geografia e Estatística (IBGE). 2014. COBERTURA E USO DA TERRA DO BRASIL 2000, 2010, 2012, 2014. ftp://geoftp.ibge.gov.br/informacoes_ambientais/cobertura_e_uso_da_terra/mudancas/vetores/. Accessed: August 2016.

Instituto Brasileiro de Geografia e Estatistica (IBGE). 2015. Mudanca na Cobertura e Uso da Terra 2000, 2010 and 2012. https://biblioteca.ibge.gov.br/index.php/biblioteca-catalogo?view=detalhes&id=294724. Accessed: August 2016

IUCN (2001) IUCN Red List categories and criteria Version 3.1. Gland, Switzerland and Cambridge, UK: IUCN Species Survival Commission

IUCN 2017. IUCN Red List of Threatened Species. Version 2017-1 www.iucnredlist.org

Iwamura, T., Possingham, H. P., Chadès, I., Minton, C., Murray, N. J., Rogers, D. I.,…Fuller, R. A. (2013). Migratory connectivity magnifies the consequences of habitat loss from sea-level rise for shorebird populations. Proceedings of the Royal Society of London B: Biological Sciences, 280, 20130325.

Joppa, L. N., O’Connor, B., Visconti, P., Smith, C., Geldmann, J., Hoffmann, M.,…&Ahmed, S. E. (2016). Filling in biodiversity threat gaps. Science, 352, 416–418.

Krauss, J., Bommarco, R., Guardiola, M., Heikkinen, R. K., Helm, A., Kuussaari, M.,…& Poyry, J. (2010). Habitat fragmentation causes immediate and time-delayed biodiversity loss at different trophic levels. Ecology letters, 13, 597–605.

Lambin, E. F., & Meyfroidt, P. (2011). Global land use change, economic globalization, and the looming land scarcity. Proceedings of the National Academy of Sciences, 108, 3465–3472.

Lockwood, J.A. (2004). Locust: the devastating rise and mysterious disappearance of the insect that shaped the American frontier. USA, Basic Books.

Mantyka-pringle, C. S., Martin, T. G., & Rhodes, J. R. (2012). Interactions between climate and habitat loss effects on biodiversity: a systematic review and meta-analysis. Global Change Biology, 18, 1239–1252.

Martinelli, G. and Moraes, M.A. (2013). Livro vermelho da flora do Brasil. Brazil, IUCN.

Moran, D., & Kanemoto, K. (2017). Identifying species threat hotspots from global supply chains. Nature Ecology & Evolution, 1, 0023.

Myers, N. (1993). Biodiversity and the precautionary principle. Ambio, 74–79.

Newbold, T., Hudson, L. N., Hill, S. L., Contu, S., Gray, C.L., Scharlemann, J.P.W.,…Purvis, A. (2015). Global effects of land use on local terrestrial biodiversity. Nature, 520, 45.

Pearson, R. G., Stanton, J. C., Shoemaker, K. T., Aiello-Lammens, M. E., Ersts, P. J.,Horning, N.,…Akgakaya, H. R. (2014). Life history and spatial traits predict extinction risk due to climate change. Nature Climate Change, 4, 217.

Pfeifer, M., Lefebvre, V., Peres, C. A., Banks-Leite, C., Wearn, O. R., Marsh, C. J.,…Cisneros, L. (2017). Creation of forest edges has a global impact on forest vertebrates. Nature, 551, 187.

Rio+20 United Nations Conference on Sustainable Development. Available: http://www.uncsd2012.org/. Accessed July 2017

Richards, P. (2015). What drives indirect land use change? How Brazil’s agriculture sector influences frontier deforestation. Annals of the Association of American Geographers, 105, 1026–1040.

Rondinini, C., Di Marco, M., Chiozza, F., Santulli, G., Baisero, D., Visconti, P.,…Amori, G. (2011). Global habitat suitability models of terrestrial mammals. Philosophical Transactions of the Royal Society B: Biological Sciences, 366, 2633–2641.

Rondinini, C., Stuart, S., & Boitani, L. (2005). Habitat suitability models and the shortfall in conservation planning for African vertebrates. Conservation Biology, 19, 1488–1497.

Drever, R. C., Drever, M. C., & Sleep, D. J.H. (2012). Understanding rarity: A review of recent conceptual advances and implications for conservation of rare species. The Forestry Chronicle, 88, 165–175.

de Ruiter, H., Macdiarmid, J. I., Matthews, R. B., Kastner, T., Lynd, L. R., & Smith, P. (2017). Total global agricultural land footprint associated with UK food supply 19862011. Global environmental change, 43, 72–81.

van Soesbergen, A., Arnell, A. P., Sassen, M., Stuch, B., Schaldach, R., Göpel, J.,…Palazzo, A. (2017). Exploring future agricultural development and biodiversity in Uganda, Rwanda and Burundi: a spatially explicit scenario-based assessment. Regional Environmental Change, 17, 1409–1420.

Sparks, T. H., Butchart, S. H., Balmford, A., Bennun, L., Stanwell-Smith, D., Walpole, M.,…Collen, B. (2011). Linked indicator sets for addressing biodiversity loss. Oryx, 45, 411419.

Strassburg, B. B., Brooks, T., Feltran-Barbieri, R., Iribarrem, A., Crouzeilles, R., Loyola, R.,…Soares-Filho, B. (2017). Moment of truth for the Cerrado hotspot. Nat. Ecol. Evol, 1, 13. The Ramsar Convention on Wetlands. Available: http://www.ramsar.org/. Accessed July 2017

Thomas, C. D., Cameron, A., Green, R. E., Bakkenes, M., Beaumont, L. J., Collingham, Y. C.,…Hughes, L. (2004). Extinction risk from climate change. Nature, 427, 145.

United Nations Millennium Development Goals. Available: http://www.un.org/millenniumgoals/. Accessed July 2017.

United States Geological Survey. 2006 Shuttle Radar Topography Mission 3 arc second version 2.0. See http://www.landcover.org/data/srtm (Accessed November 2016).

Veach, V., Di Minin, E., Pouzols, F. M., & Moilanen, A. (2017). Species richness as criterion for global conservation area placement leads to large losses in coverage of biodiversity. Diversity and Distributions, 23, 715–726.

Visconti, P., Pressey, R. L., Giorgini, D., Maiorano, L., Bakkenes, M., Boitani, L.,…Rondinini, C. (2011). Future hotspots of terrestrial mammal loss. Philosophical Transactions of the Royal Society of London B: Biological Sciences, 366, 2693–2702.

Visconti, P., Bakkenes, M., Baisero, D., Brooks, T., Butchart, S. H., Joppa, L.,…Maiorano, L. (2016). Projecting global biodiversity indicators under future development scenarios. Conservation Letters, 9, 5–13.

World Wildlife Fund, 2015. The Big Five of the Cerrado. http://www.wwf.org.br/informacoes/english/750242/The-Big-Five-of-the-Cerrado, Accessed August 2016.

